# Developmental programmes drive cellular plasticity, disease progression and therapy resistance in lung adenocarcinoma

**DOI:** 10.1101/2024.12.03.626626

**Authors:** Kamila J Bienkowska, Stephany Gallardo Y, Nur S Zainal, Matthew Ellis, Maria-Antoinette Lopez, Judith Austine, Sai Pittla, Serena J Chee, Aiman Alzetani, Emily C Shaw, Christian H Ottensmeier, Gareth J Thomas, Christopher J Hanley

## Abstract

**Background:** Cellular plasticity, involving loss of lineage determination and emergence of hybrid cell states, plays a pivotal role in non-small cell lung cancer (NSCLC) disease progression and therapy resistance. However, the full spectrum of atypical states generated in human NSCLC and the pathways that regulate them are yet to be fully elucidated. Here we examine the role of developmental programmes, alveogenesis and branching morphogenesis (BM), in regulating phenotypic diversity in NSCLC.

**Methods:** Transcriptomic analysis of epithelial cells isolated from murine lungs at different stages of organogenesis were used to derive gene signatures for developmental programmes. Bulk tissue transcriptomic datasets from human NSCLC and non-neoplastic control samples were used to identify whether developmental programmes were associated with molecular, morphological, and clinical parameters. Single-cell RNA-sequencing was used to identify malignant cell states in human NSCLC (n = 16,621 epithelial cells from 72 samples) and protein level validation of these states was carried out using multiplexed immunohistochemistry (n = 40).

**Results:** Mutually antagonistic regulation of alveogenesis and BM was found to account for a significant proportion of transcriptomic variance in human NSCLC bulk tissue datasets. BM activation was associated with poor overall survival rates in five independent lung adenocarcinoma (LUAD) cohorts (p=2.04e-13); and was significantly prognostic for resistance to tyrosine kinase inhibitors (TKIs; p=0.003) and immune checkpoint blockade (ICBs; p=0.014), in pre-treatment biopsies. Single-cell RNA-sequencing analysis revealed that malignant LUAD cells with loss of alveolar lineage fidelity predominantly acquired inflamed or basal-like cellular states, which were variably persistent in samples from TKI and ICB recurrence.

**Conclusions:** Our results show LUAD tumours undergo reversion from an alveogenic to branching morphogenic phenotype during disease progression, generating inflamed or basal-like cell states that are variably persistent following TKI or ICB treatments. These findings identify prognostic biomarkers for therapy response and underscore the role of different cell states in resistance to multiple treatment modalities.

## Introduction

Lung cancer is the leading cause of cancer-related deaths worldwide [1], despite recent advances in basic research and the implementation of oncogene- or immune-targeted therapies [2]. Non-small cell lung cancers (NSCLC) represent 85% of all lung cancer cases, which are subdivided into two main subtypes: lung adenocarcinoma (LUAD) and lung squamous cell carcinoma (LUSC) [3]. LUAD is the most common of these subtypes and the predominant type of lung cancer found in never-smokers [3]. LUAD tumours typically develop in peripheral regions of the lung, mainly arising from oncogenic transformation of alveolar type 2 cells (AT2) [4]. LUSC originates from squamous airway epithelial cells predominantly found in more central regions of the lung [4].

Cell turnover in healthy adult lungs is low, but following injury, progenitor populations can become activated to proliferate and differentiate into one or more cell types [5,6]. This increased epithelial plasticity has also been described in LUAD primary tumours, leading to loss of lineage fidelity and the emergence of mixed-lineage or regenerative phenotypes [5,7]. Furthermore, plasticity has also been associated with metastatic progression and histological subtype determination, involving transcription factors that regulate embryonic development (SOX2 and SOX9) [5,8].

Normal lung development involves two independent and mutually antagonistic programmes: branching morphogenesis (BM) and alveogenesis (ALV) [9]. BM describes the process of coordinated proliferation and collective migration leading to development of the airways and the proximal-distal axis [10]. ALV involves the generation and maturation of alveoli, enabling efficient gas exchange [10].

In this study, we performed a comprehensive analysis of transcriptomic (bulk-tissue and single cell) datasets to determine whether aberrant regulation of developmental programmes (ALV and BM) influence human NSCLC disease progression and therapy resistance.

## Methods

### ALV and BM gene signatures

Data from Chang *et al.* [9] was used to generate ALV/BM gene signatures. The top 100 upregulated genes (ranked by log2 Fold Change) on embryonic day 14 (active branching morphogenesis) and embryonic day 19 (active alveogenesis) were used to generate the provisional BM and ALV signatures, respectively (**Table S1**). Significantly enriched (Bonferroni adjusted p value < 0.05) Gene-ontology (GO) terms were identified using the *enrichGO* function from the ClusterProfiler R package [11], setting the background gene set to all genes reported in the Chang *et al.* [9] study.

71 ALV genes were present in TCGA data, and 76 in scRNAseq data. 82 BM genes were present in both TCGA and scRNAseq data. Single-sample Gene Set Enrichment Analysis (ssGSEA) [12] implemented in the GSVA (Gene Set Enrichment Variation) R package [13] was used to calculate enrichment scores of ALV and BM gene signatures per sample: Gene signatures (lists) of interest were set as “gset.idx.list” argument, while a gene expression matrix containing all genes in the analysed dataset was set as “expr” argument. The “method” argument was set to “ssgsea”.

### Bulk tissue transcriptomic data processing and analysis

#### Data access and processing

NSCLC TCGA RNA-seq dataset (Firehose Legacy) was downloaded into RStudio using TCGAbiolinks package [14] and processed using the variance stabilising transformation implemented in DE-Seq2 R package [15]. Raw RNA-sequencing data from GSE103584 [16] was downloaded and processed to logarithmically normalised counts per million (lcpm) using recount3 R package [17]. Genes with few or no counts across samples were removed from the dataset using *filterByexprs* function (edgeR package [18]) with default parameters, and argument “group” set to “Histopatholoy”. TMM (trimmed mean of M-values) method was used to determine normalisation factors, and the data was prepared in the cpm (counts per million) format, then log-transformed (lcpm) and scaled. Microarray datasets (GSE31552 [19], GSE72094 [20], GSE31210[21], GSE68465 [22], GSE157009 [23], GSE17010 [23] and GSE4573 [24]) were log-transformed (if not already performed prior) to obtain normal distribution and quantile-normalised using NormalizeBetweenArrays function (limma package [25]). mRNA expression level z-scores were downloaded for the IMPACT study cohort, accessed through the OncoSG Cancer Genomics Data portal (https://src.gisapps.org/OncoSG/). Bulk tissue RNA-sequencing data from ICB datasets were downloaded from GSE207422 and GSE135222 using the pre-processed transcript per million (TPM) values, which were transformed to log2(TPM+1) for analysis.

#### Principal Component Analysis (PCA)

PCA was implemented using the *prcomp* function (base R). The results were visualised with the use of factoextra [26] package. The contribution of ALV/BM genes to each PC was assessed using a Fisher’s exact test to measure enrichment amongst the top quartile of genes with the highest absolute loading coefficients.

#### Survival analysis

Survival analysis was performed using the survival and survminer R packages. The *surv_categorize* function was used to group samples into BM_high and BM_low groups, based on ssGSEA scores. The cut-point value used for stratification was calculated using the maximally selected rank statistics from the maxstat R package. 5-year overall survival (OS) or progression free survival was used as the experimental outcome as indicated in the relevant Figures. Kaplan-Meier plots were prepared using *ggsurvplot* and Log-rank tests were performed to calculate p values. Cox proportional hazard regression model (*coxph* function) was fitted based on 5-year overall survival data and with stage, age, and BM expression levels as covariates. Hazard ratios with 95% confidence intervals were visualised as a forest plot.

### Single-cell RNAseq data processing and analysis

#### Data processing

Four single-cell RNA-sequencing datasets of NSCLC were analysed in this study: “Kim” [27], “Czbiohub” [28], “Zilionis” [29] and TLDS (Target Lung Dataset) [30]. Only samples originating from primary lung tumours or normal tumour-adjacent tissues were used to create the integrated dataset.

We used the Travaglini *et al.* [31] Human Lung Cell Atlas as a reference dataset to train a machine learning classifier to identify cell-types within human lung tissues (Seurat’s Label Transfer). Single cells were assigned to appropriate cell types (meta lineages) if a prediction probability for this cell type was greater than 0.75.

Cells assigned as epithelial cells were extracted from each dataset and integrated using the Seurat R package [32] to implement the method described by Stuart *et al.* [33]. This was achieved, first by defining anchors connecting the datasets (using the *FindIntegrationAnchors* function) based on 30 principal components (PCs) and 2000 variable features. *The IntegrateData* function was run to integrate the datasets and correct for batch-effects.

PCA was then performed on the scaled dataset. *FindNeighbors* and *FindClusters* functions were used to cluster cells, and *FindAllMarkers* function was utilised to calculate marker genes defining these clusters (Wilcoxon Sum Rank test, log2FC ≥. 0.25, min.pct = 0.1) Any mis-labelled cells (by the machine learning classifier) tended to be cell doublets (expressing genes typical to more than one cell lineage) and were filtered out.

#### Module scores

Seurat’s *AddModuleScore* function was used to calculate single-cell expression levels for WGCNA modules and developmental programme gene lists. For WGCNA modules all genes assigned to the relevant module with a positive module membership score (MM > 0) were used to calculate these scores for epithelial cells within the integrated scRNA-seq dataset. Module scores were calculated for the developmental programmes on epithelial cells isolated from the complete (including metastatic sites) Kim [27], Maynard [28] and GSE207422 [34] datasets following log-normalisation.

#### Epithelial Cluster identification and differential gene expression analysis

Cluster identification was performed on the “Integrated” assay using Seurat’s *FindClusters* function (resolution was set to 0.05). Gene markers of cell populations were identified with the use of Seurat’s *FindConservedMarkers* function run on the “RNA” assay (Wilcoxon test; log2FoldChange ≥ 0.1, genes expressed in at least 10% of cells in either group, grouping variable set to Dataset). Genes were considered statistically significant where the adjusted p <0.05 in all datasets analysed. For analysis of all cells across epithelial subpopulations all four datasets were analysed. For analyses comparing LUAD-basal to LUAD-Inflamed cells three datasets were analysed due to low cell numbers (<100 cells) for these clusters in the TLDS dataset.

#### AT2 and Basal cell prediction score (Lineage Fidelity)

Label transfer analysis, as described by Stuart *et al.* [33], was used to calculate basal and AT2 prediction scores using Travaglini’s [31] population of normal basal epithelial cells or AT2 cells as a reference, respectively. First, *FindTransferAnchors* function was used to calculate anchors between the reference population and the integrated scRNAseq epithelial cells. Then, the *Transferdata* function returned basal/AT2 prediction scores for every queried cell (values ranging from 0 to 1).

### Mutational signatures and TP53/KRAS mutations

TCGA data containing mutational information (LUAD/LUSC, Firehose Legacy) were downloaded into R and converted into maf objects using TCGAmutations and maftools [35] packages. LUAD samples were stratified into BM_high and BM_low groups on the cut-point value from the survival analysis (ssGSEA score = 0.1241903). To extract mutational signatures present in the datasets, we first used the *trinucleotideMatrix* function (NFM package [36]) to extract single 5’ and 3’ bases flanking mutated sites, followed by *extractSignatures* that allowed the signature identification. In order to calculate differential enrichment of a signature across LUAD groups, we used the contribution of each signature to the overall mutational profile of a sample (sum of all signatures = 1).

### Histology staining

#### Tissue microarray (TMA) construction

Archival formalin-fixed paraffin-embedded (FFPE) material was used to construct a TMA from 1-mm cores (Aphelys Minicore 2, Mitogen). Regions for inclusion in TMA cores were selected by a pathologist (E. C. S.) to include areas of well-differentiated and poorly differentiated (solid) LUAD (n= 30 cores from 10 patients for each), LUSC (n= 30 cores from 10 patients) and tumour adjacent non-neoplastic lung tissue (n= 30 cores from 10 patients). Ethical approval was obtained through the UK National Research Ethics Service (NRES; Rec No. 10/H0504/32), and informed consent was obtained from each patient. All tissue collection and storage were handled by the Department of Cellular Pathology at University Hospital Southampton NHS Trust.

#### Multiplexed immunohistochemistry and Histo-cytometry analysis

Automated immunostaining (PT Link Autostainer, Dako) was performed in a Clinical Pathology Accreditation (UK) Ltd (CPA)-accredited clinical cellular pathology department. Cyclical multiplexed immunohistochemistry was performed as described previously [37]. Briefly, serial four-micron tissue sections from a TMA block were stained using the antibodies detailed in **Table S2.** CD31 staining was carried out using DAB substrate and used as fiducial points for image registration across imaging iterations. Subsequent stains were performed using AEC substrate to enable clearing between markers. Colour deconvoluted images were generated for each antibody stain representing blue (nuclear; haematoxylin), brown (CD31; DAB) and red (panel markers; AEC) signals. Target specific staining was thresholded by identifying topological maxima above the level of background signal, measured as the moving average over a 151D×D151-pixel window (Original-Background <5D=D0). For the red signal, an additional condition was applied to account for the similarity in the colour profile of red and brown (red signalD<D(0.43*brown signal)D=D0). Quality control (QC) checks for tissue integrity throughout cyclical staining were performed on the nuclear stain, to identify tissue loss or registration errors: nuclear signal was identified from the first staining iteration (i[1]), then any nucleus with a signal missing in the final staining iteration (i[n]) were filtered out.

Histo-cytometry analysis was performed using the filtered nuclear stain to segment individual objects. These regions were then grown up to 5.5 µm radius to represent simulated cell regions. Histo-cytometry estimates of cellular expression levels were calculated as the fraction of pixels within simulated cell regions positive for staining once background levels had been subtracted. Pan-CK+ cells were identified where the expression level exceeded 0.2. For all other markers cells were classified as positive where the expression level exceeded 0.2 or the median +1*mad (whichever value was higher). Cell phenotypes were identified by positivity for one or more relevant markers and negativity for alternative phenotype markers. Cells exhibiting positivity for markers of two separate phenotypes were classified as undetermined. Subpopulation enrichment was determined using Pearson’s residuals from a chi square test assessing the number of each subpopulation as a proportion of the total number of epithelial cells across all three cores from each individual patient sample.

### Statistical Analysis

All statistical tests carried out were two-sided as specified in the relevant Figure legends. All Boxplots are displayed using the Tukey method (centre line, median; box limits, upper and lower quartiles; whiskers, last point within a 1.5x interquartile range) as implemented in the *geom_boxplot* function from the ggplot2 R package. Unless otherwise specified, statistical significance based on group comparisons was performed using the *geom_pwc* function from ggpubr package. Details on the exact tests used are included in Figure captions. Where asterisks are used to represent statistical significance, these represent the following p-values: Ns: p > 0.05. *: p ≤ 0.05, **: p ≤ 0.01, ***: p ≤ 0.001, ****: p ≤ 0.0001.

## Results

### Developmental gene expression programmes are a key source of transcriptomic variance in NSCLC

We hypothesised that NSCLC plasticity was driven by the aberrant regulation of developmental programmes (BM and ALV) (**Figure 1A**). To test this hypothesis, we first used a publicly available microarray dataset [9] to identify genes differentially expressed between epithelial cells engaged in BM (embryonic day 14 [E14]) or ALV (embryonic day 19 [E19]) (**Figure 1B and TABLE S1**). Functional annotation showed that the BM signature was significantly enriched in mitotic processes (**Figure 1C**). In contrast, the ALV signature was associated with canonical AT2 cell functions, such as surfactant homeostasis, in addition to regulation of inflammatory responses, body fluid levels, and angiogenesis [38] (**Figure 1D**).

**Figure 1.**
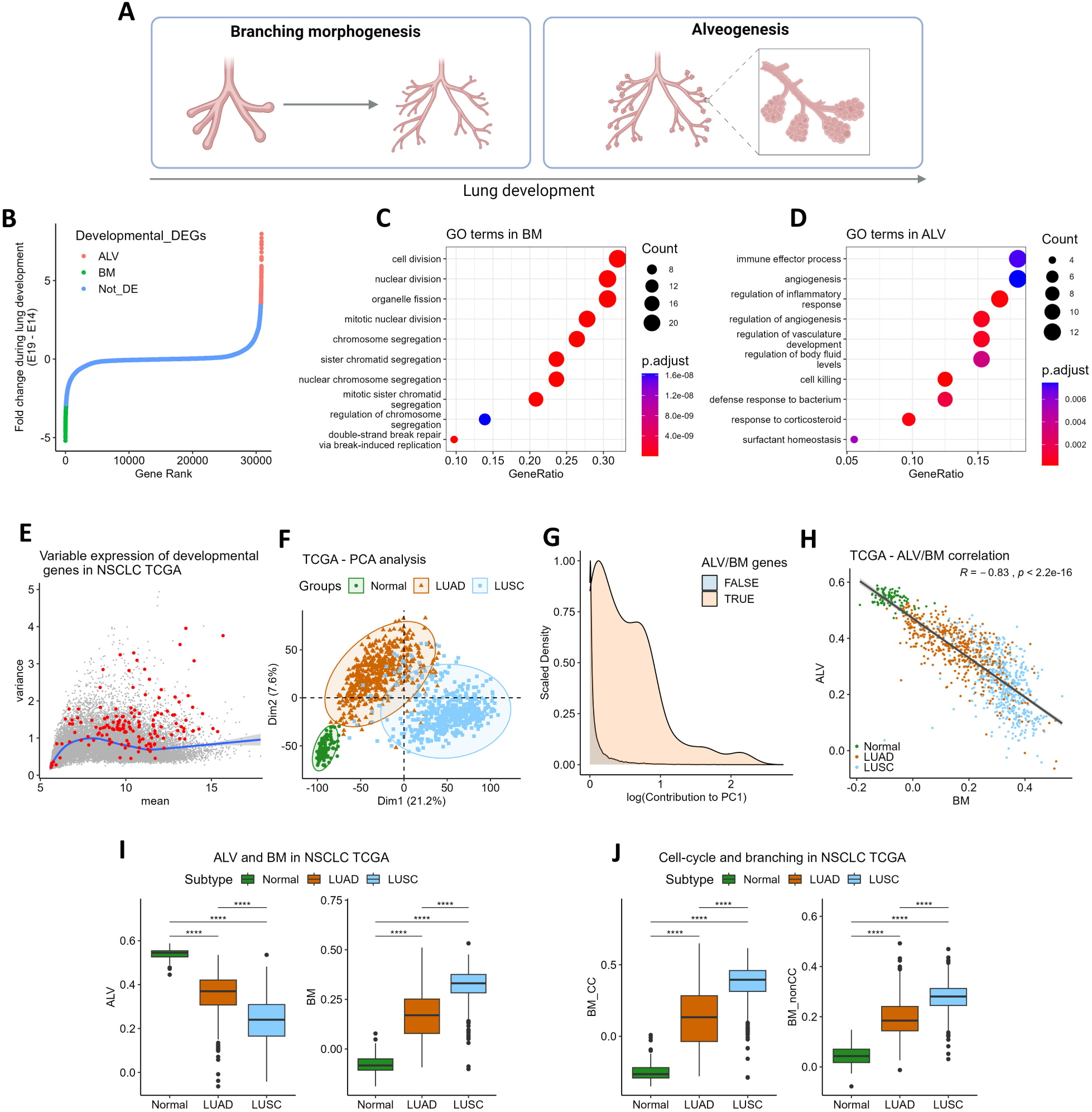
Developmental Alveogenesis (ALV) and Branching Morphogenesis (BM) programmes are associated with transcriptomic variance in NSCLC. **A** – Schematic showing the different stages of lung development. Created with Biorender.com. **B** – Dot plot showing the identification of genes differentially expressed by epithelial cells during murine developmental-BM (embryonic day 14, green) and ALV (embryonic day 19, red) [9]. **C,D** – Dot plots showing GO terms enriched in ALV/BM genes. **E** – Scatter plot showing mean expression level and variance for all genes in NSCLC samples from TCGA (RNA-Seq); BM/ALV genes shown in red. **F** – PCA of NSCLC samples. **G** – Histogram showing the contribution of ALV/BM genes to PC1. **H** – Scatter plot showing ALV and BM scores across NSCLC samples; *R* = Spearman’s correlation. **I,J** – Boxplots showing enrichment scores in TCGA cohort grouped by histological subtype, calculated using ssGSEA: for ALV and BM signatures (I); and BM_cc (cell-cycle related genes) and BM_nonCC (non-cell-cycle related genes) (J). Statistical significance was assessed by Wilcoxon test for pairwise comparisons (asterisks represent Bonferroni adjusted p-values).

We then analysed the expression of ALV and BM signatures in NSCLC, using The Cancer Genome Atlas (TCGA) RNA-seq dataset, consisting of LUAD (n = 517), LUSC (n = 501), and control (non-neoplastic) lung tissue samples (n = 110). ALV and BM genes exhibited high expression-level standardised variance, implying an important role in NSCLC biology (**Figure 1E**). PCA (Principal Component Analysis) showed components 1 and 2 separated the NSCLC samples by subtype (**Figure 1F**) and genes from the ALV/BM signatures heavily influenced this separation (**Figure 1G & S1A-B**). We also observed a negative correlation between enrichment scores for the ALV/BM signatures in NSCLC samples (**Figure 1H**, r = - 0.83, p<2.2e-16), consistent with their antagonistic roles in lung development [9]. The ALV signature was significantly downregulated in both tumour subtypes compared to control samples, whereas BM was upregulated. Furthermore, LUSC samples had significantly lower enrichment scores for ALV and higher scores for BM than LUAD (**Figure 1I**).

Cell proliferation is fundamental to both developmental BM and cancer. To determine whether the association between BM and cancer extended beyond genes involved in this process we split the BM signature into genes associated with cell cycle, cell proliferation, cell division and cell replication gene ontology terms (BM_CC); and non-proliferation associated genes (BM_nonCC), comparing enrichment scores for these new signatures across NSCLC samples. This showed both BM_CC and BM_nonCC signatures varied similarly between NSCLC subtypes (**Figure 1J**).

These findings were validated using a Laser Capture Micro-Dissected (LCMD) microarray dataset [19]: comparing the epithelial compartment from non-tumour alveoli (NTa), not-tumour bronchi (NTb), LUAD and LUSC (**Figure S1C-J**). Additionally, we confirmed the negative correlation between ALV and BM signatures in multiple (LUAD [n = 3] and LUSC [n = 3]) independent microarray datasets (**Figure S1K**) [20–24].

Given the variation in ALV/BM gene expression observed between NSCLC subtypes and non-neoplastic alveolar/bronchial samples, we examined whether the tumour’s location along the proximal-distal axis influenced ALV/BM signature expression in NSCLC. This showed an increase in ALV and decrease in BM for peripheral tumours compared to central tumours (**Figure S1L**). However, ALV/BM scores were not significantly different when comparing central and peripheral tumours from LUAD or LUSC subtypes independently (**Figure S1L**).

In summary, these analyses demonstrated that ALV and BM gene expression programmes strongly influenced transcriptomic variation in NSCLC. LUSC uniformly suppressed ALV and activated BM; whereas LUAD tumours were heterogeneous for BM activation with some cases exhibiting ALV programme expression comparable to control tissue.

### BM activation is associated with poor outcome in lung adenocarcinomas

To determine the clinical impact of BM activation we investigated whether this gene expression programme was associated with survival rates in NSCLC patients, using nine independent datasets (n = 5 LUAD and 4 LUSC). This showed that the BM signature identified LUAD patients with significantly reduced five-year overall survival rates in each cohort analysed (TCGA, GSE72094 [20], GSE31210 [21], GSE68465 [22], GSE103584 [16]; n = 1680 patients in total; **Figure 2A and S2A**). Multivariate analysis showed that BM activation was a significant prognostic factor independent of disease stage and age (HR [95% CIs] = 2.12 [1.73-2.59], p = 2.04e-13; **Figure 2B**). In contrast, there was no consistent association with survival in LUSC cohorts (TCGA, GSE157009 [23], GSE157010 [23], GSE4573 [24] (**Figure S2B**).

**Figure 2.**
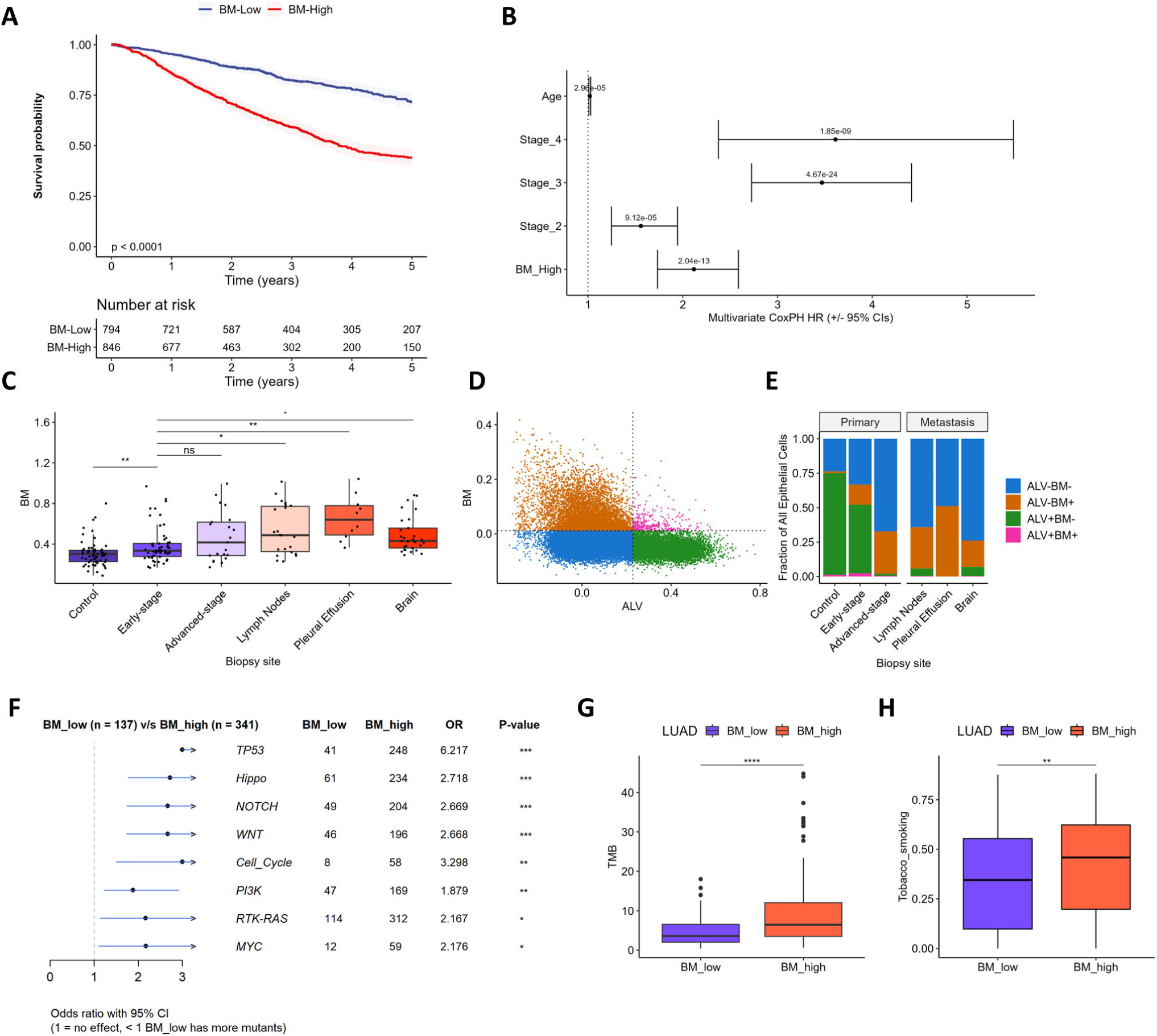
High expression of BM in LUAD predicts poor survival and is associated with frequent TP53 mutations. A,B. – 5-year overall survival (OS) analysis in LUAD stratified by BM expression across 5 combined LUAD datasets (TCGA, GSE72094 [20], GSE68465 [22], GSE31210 [21], GSE103584 [21]): (**A**) Kaplan-Meier (KM) plot; (**B**) Forest plot showing multivariate Cox proportional hazards regression model coefficients (+/− 95% CIs) and adjusted p-values. **C-E –** BM expression in advanced and metastatic LUAD samples (from Kim *et al.* [27]): (**C**) Boxplots showing sample-level epithelial BM ssGSEA scores; (**D**) Scatter plot showing classification of epithelial cells based on ALV/BM expression, dotted lines represent signature positivity thresholds (median + 1 median absolute deviation); (**E**) Bar plot showing proportion of epithelial cells defined by ALV/BM classes in primary and metastatic LUAD. **F** - Forest plot showing the Odds ratio (OR) and Fischer’s exact test P-values for genetic perturbations in oncogenic pathways comparing BM-high vs BM-low LUAD samples. **G,H** – Boxplots showing tumour mutational burden (TMB, **G**) and tobacco smoking COSMIC mutational signature score (**H**) in LUAD samples stratified by BM expression. Wilcoxon test P values are shown in C, G, H.

We next investigated whether BM activation increased during disease progression. For this analysis we utilised the Kim *et al.* dataset [27] where scRNA-seq was performed on samples taken from different disease stages and anatomical sites. This showed that BM activation was significantly increased in epithelial cells from both local and distant LUAD metastases (**Figure 2C**). Furthermore, analysis at the single-cell level showed that ALV-/BM- and ALV-/BM+ LUAD cells were increased in both advanced stage primary tumours and metastases (**Figure 2D,E**).

We then examined whether there were any genetic changes associated with BM activation. This showed that *TP53* (65%) and *KRAS* (37%) were the most frequently mutated genes in LUAD BM-high and LUAD BM-low samples, respectively (**Figure S2C**). Furthermore, the odds of TP53 signalling pathway being altered were significantly higher in the BM-high group (Odds ratio = 6.217, 95% CI, 6.22-9.92, adj.p = 7.9e-17) (**Figure 2F**). Interestingly, LUAD samples containing *TP53* mutations but not *KRAS* mutations showed the highest BM scores (**Figure S2D**).

BM-high LUAD tumours were also found to have significantly higher tumour mutational burden (TMB) (**Figure 2G**), which is known to be associated with smoking [39]. Consistent with this, the tobacco smoking mutational COSMIC signature (SBS4) was significantly enriched in the BM-high compared to BM-low LUAD tumours (**Figure 2H**).

In summary, we showed that BM activation in LUAD is associated with disease progression and poor survival. LUAD-BM tumours were also characterised by TP53 pathway mutations, increased TMB and enrichment for the tobacco smoking mutational signature.

### BM activation is associated with targeted-therapy resistance in lung adenocarcinomas

Loss of alveolar features has been linked to tyrosine kinase inhibitor (TKI) resistance [28,40]. Given the inverse relationship between ALV and BM we suspected that BM activation may also be involved in TKI resistance. To investigate this, we first analysed the Maynard *et al.* dataset [28] (**Figure 3A**), where longitudinal scRNA-seq analysis was performed on samples collected from patients undergoing a range of TKI treatments (**Figure S3A,B**). This showed low levels of BM activation in tumour cells from residual disease (RD) that was significantly increased in samples with recurrent progressive disease (PD) (**Figure 3B**). These changes were not associated with increased stage in recurrent samples as a similar increase was observed when only stage IV cases were analysed (**Figure S3C**). Single-cell analysis showed that both ALV-BM- and ALV-BM+ LUAD cells were increased in samples from recurrent progressive disease (**Figure 3C,D**).

**Figure 3.**
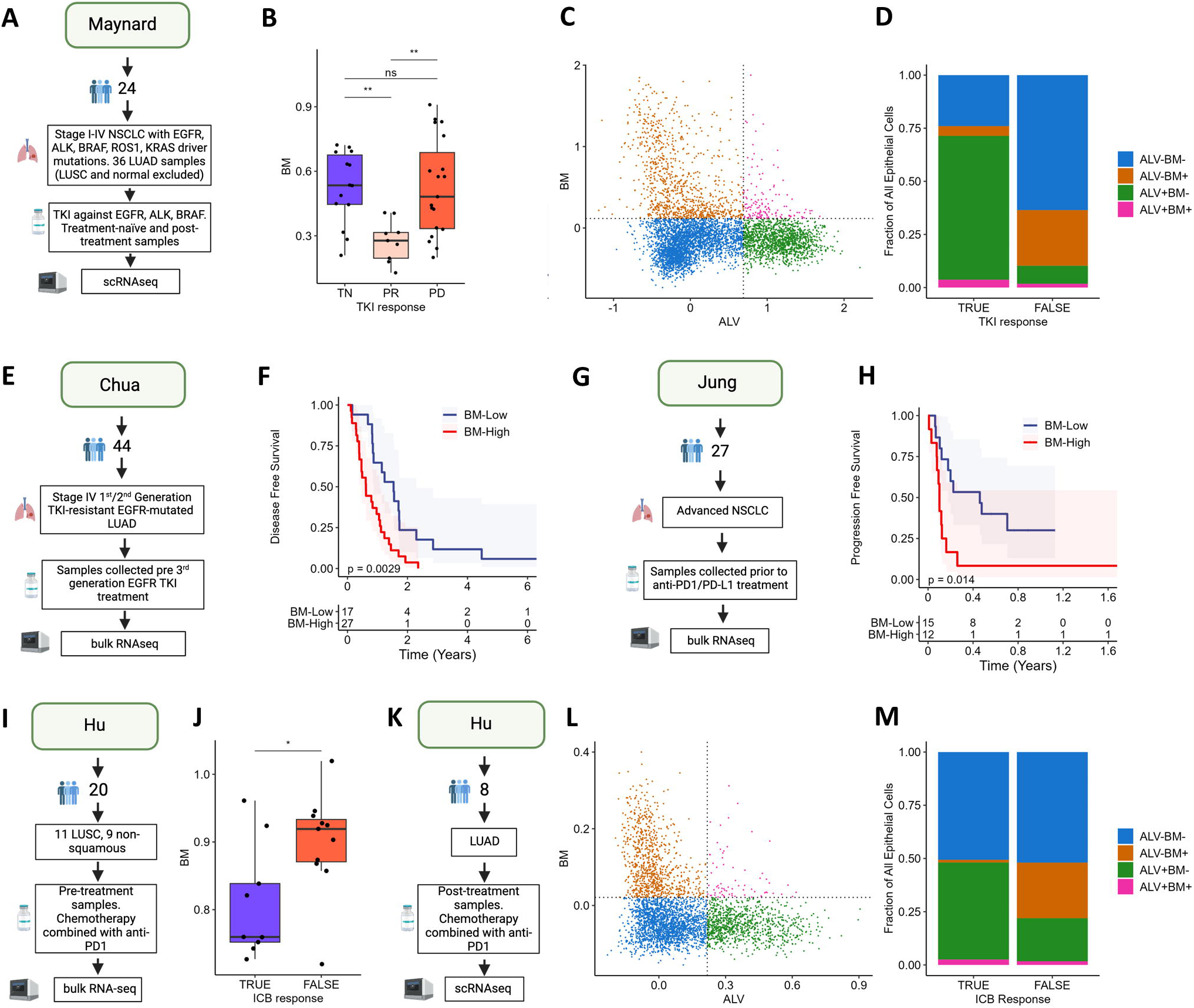
High expression of BM is associated with resistance to therapies. **A,E,G,I,K**– Schematics illustrating the designs of the analysed datasets. **B** – BM expression in TKI treated LUAD samples (from Maynard *et al.* [28]); TN – treatment naïve, PR – partial response, PD – progressive disease). **C -** Scatter plot showing classification of epithelial cells based on ALV/BM expression (Maynard), dotted lines represent signature positivity thresholds (median + 1 median absolute deviation). **D** - Bar plot showing proportion of epithelial cells defined by ALV/BM classes in LUAD samples grouped by response to TKI (Maynard). **F** - KM plot showing disease-free survival of patients receiving EGFR inhibitors (IMPACT study; from Chua *et al.* [40]) stratified based on pre-treatment BM expression; **H –** KM plot showing progression-free survival of patients with advanced NSCLC receiving anti-PD-1/PD-L1 (from Jung *et al.* [41]) grouped by BM expression. **J** - BM expression in NSCLC patients receiving chemotherapy treatment combined with anti-PD1, grouped by response to treatment (from Hu *et al.* [34]); TRUE/FALSE refers to major pathologic response). **L,M –** as per panels C,D. Log-rank test P values are shown in F and H. Wilcoxon test P values are shown in B, J.

EGFR mutations are the most common actionable drivers in LUAD and resistance to 1^st^ or 2^nd^ generation EGFR inhibitors (EGFRi) is commonly attributed to T790M alterations, which can now be targeted by 3^rd^ generation inhibitors. The IMPACT study examined the efficacy of 3^rd^ generation EGFRi in a cohort of 1^st^/2^nd^ generation EGFRi resistant stage IV LUAD patients with mixed T790M status [40] (**Figure 3E**). BM activation was increased in samples lacking the T790M alteration (**Figure S3D)**, consistent with the loss of alveolar lineage fidelity in these samples described previously. BM activation was also found to be a significant prognostic indicator in this cohort (n = 44, p 0.0029; **Figure 3F**). Notably, BM activation was more effective in stratifying patient response rates than T790M status (**Figure S3E**). When combining the two variables in a multivariate cox regression model BM activation was the only independently significant predictor of survival rates and in cases with T790M mutations survival rates could also be significantly stratified by BM status (**Figure S3F,G**).

Previous studies have shown that high TMB and *TP53* alterations can limit the efficacy of chemotherapy and TKI treatments [42], which may in part explain the association between BM and TKI resistance. However, these genetic features have also been identified as favourable prognostic indicators for immune checkpoint blockade (ICB) [43]. Moreover, using RPPA data from TCGA cohort we found that PD-L1 was the third most significantly up-regulated protein in BM-high vs BM-low LUAD samples (**Figure S3H**). Therefore, three key biomarkers for favourable ICB response [44] were upregulated in BM-high LUAD. This suggested that while BM-high LUAD tumours were refractory to treatments directly targeting the tumour cells, they may be more vulnerable to ICB. To test this hypothesis, we analysed BM activation in pre-treatment biopsies from patients that received ICB monotherapy [41] (**Figure 3G**). Contrary to expectations, we found BM-high tumours had significantly reduced progression free survival rates (**Figure 3H**). Further analysis of a second cohort [34] (**Figure 3I**) confirmed this finding, showing BM activation was significantly reduced in tumours that showed a major pathological response to combined ICB and chemotherapy treatment (**Figure 3J)**. We analysed scRNA-seq data from post-treatment LUAD samples in the Hu *et al.* dataset [34] (**Figure 3K**) to determine whether ICB treatment resulted in changes in BM activation as we had found following TKI treatment. This showed that the ALV-BM+ LUAD population was enriched in post-treatment samples with no major pathological response to combined ICB and chemotherapy treatment, but notably (and unlike TKI resistant samples) the proportion of ALV-BM-LUAD cells was unchanged (**Figure 3L,M**).

In summary, BM activation was a key factor in promoting LUAD progression and resistance to current treatments identifying patients that, although positive for ICB response biomarkers, will likely fail to respond to this treatment.

### BM activation in LUAD is associated with loss of alveolar lineage determination

To investigate ALV and BM activation in NSCLC further, we integrated four scRNA-seq NSCLC datasets: “Kim” [27], “Czbiohub” [28], “Zilionis” [29] and TLDS (Target Lung Dataset) [37]. We then performed cell-type classification using a machine learning classifier, trained on the Human Lung Cell Atlas (HLCA) described by Travaglini *et al.* [31] (**Figure 4A**). Cells classified as epithelial were extracted from each dataset, integrated to mitigate batch effects [33], and manually filtered to exclude doublets. This generated a dataset containing 16,621 epithelial cells from 72 samples: 69% LUAD (n = 42); 7.5% LUSC (n = 13) and 23.5% control (non-neoplastic; n = 17). Transcriptomic variation between control, LUAD and LUSC epithelium was clear following UMAP visualisation (**Figure 4B**). Consistent with the analyses presented above, pseudo bulk expression profiles for each sample showed that ALV and BM scores were significantly negatively correlated (r = −0.68, p = 4.1e-09).

**Figure 4.**
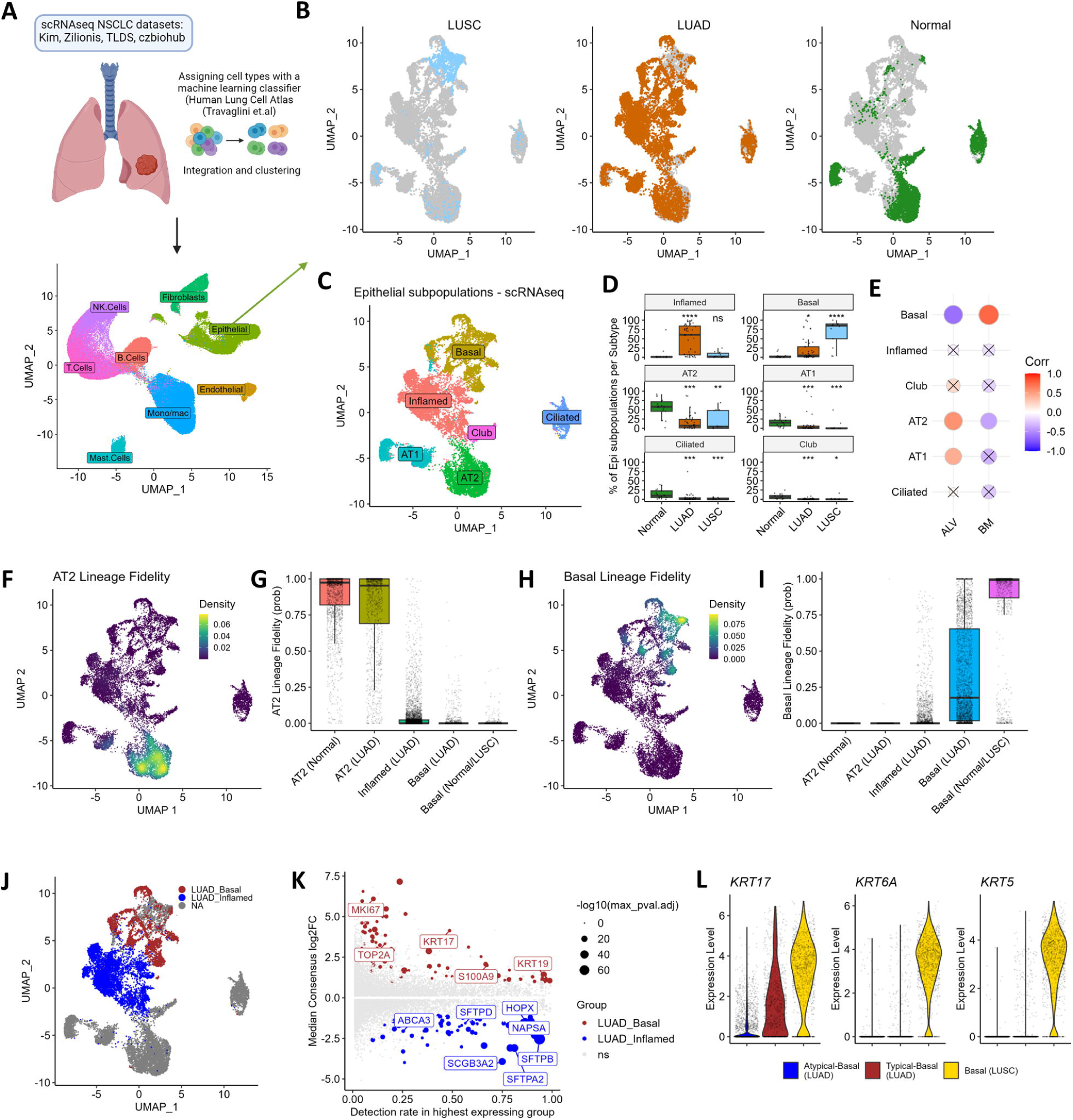
BM upregulation in LUAD is associated with acquisition of basal-like features. A,B –. NSCLC epithelial subpopulation analysis by scRNA-seq, (A) UMAP visualisation of subpopulation clustering and (B) boxplots showing the relative abundance of each subpopulation across LUAD, LUSC and normal samples. **C** – UMAP plot showing identified epithelial subpopulations. **D** – Boxplot showing the relative abundance of each subpopulation across LUAD, LUSC, and normal samples. **E** - Dot plot showing Pearson’s correlation between subpopulation abundance and ssGSEA scores for ALV/BM (p-values>0.01 are crossed out). **F,G** – UMAP (F) and Boxplots (G) showing AT2 lineage fidelity scores. **H,I** – UMAP (H) and Boxplots (I) showing Basal lineage fidelity scores. **I** – UMAP plot showing epithelial cells from LUAD samples assigned to either the Inflamed or Basal population. **J** – UMAP showing epithelial cells classified as LUAD_Inflamed and LUAD_Basal. **K**– Scatter plot showing consensus differential expression analysis (across three independent scRNA-seq datasets) comparing Inflamed and Basal subpopulations from LUAD samples, significance was determined by maximum adjusted p-value<0.05. **L** – Violin plots showing expression levels for basal keratins in LUAD cells from the Basal cluster, grouped based on their Basal Lineage Fidelity (atypical-basal < 0.5; typical-basal > 0.5) or LUSC basal cells.

Given that the epithelial compartment of adult lung tissue consists of multiple lineages, we investigated whether specific subpopulations were associated with BM activation. Unsupervised clustering identified 6 epithelial subpopulations, 5 of which represent well-described lineages: “Basal”, “AT2”, “AT1”, “Ciliated”, and “Club” (**Figure 4C**).

The “Basal” subpopulation was characterised by high molecular weight keratins (*KRT5/6B/10/14/15/16/17)* and other canonical markers (*e.g. DAPL1* and *TP63*), in addition to proliferation-associated genes (*e.g. MKI67*, *STMN1*, *TUBB, TOP2A*) (**Figure S4A**). This subpopulation was significantly enriched in cancer samples, particularly in LUSC (**Figure 4D**). Notably, 82% of all LUSC cells were assigned to the “Basal” cluster - consistent with previous studies describing basal cells as the cell-of-origin for LUSC tumours [45].

The “AT2” subpopulation was identified by overexpression of canonical markers (*e.g., SFTPC, SFTPD, SFTPA1/2, ABCA3,* and *NAPSA;* **Figure S4A**). Despite AT2 cells being well described as the cell of origin for LUAD, this population was significantly less abundant in LUAD samples compared to control, demonstrating a high degree of transcriptomic plasticity within LUAD epithelium (**Figure 4D**).

“AT1” cells were identified by overexpression of canonical markers such as *AGER, CAV1* and *EMP2* (**Figure S4A**). Interestingly, these cells formed two discrete groups in the UMAP projection (**Figure 4C**). One subset was only present in LUAD samples (**Figure 4B**) and expressed canonical AT1 marker genes at a lower level than the AT1 cells isolated from control tissue samples **(Figure S4A)**. It also expressed markers of AT2 cells, such as *NAPSA* and *SFTPD,* likely reflecting the mixed-lineage hybrid state described in previous LUAD studies [5,46,47].

Ciliated cells were identified by overexpression of *TPPP3, SNTN, CAPS, TSPAN1, CDHR3* (**Figure S4A**); and Club cells by overexpression of *SCGB1A1, SCGB3A1* and *BPIFB1* (**Figure S4A**). Both subpopulations were significantly downregulated in LUAD samples compared to control samples (**Figure 4D**).

We named the sixth population “Inflamed”. It was marked by genes also associated with prostaglandin and interferon signalling (*e.g., HPGD, IRF7*, *EREG*, *BST2*; **Figure S4A**). This population was the most abundant phenotype in LUAD samples and significantly enriched in LUAD samples compared to control samples (**Figure 4D**). Notably, this population expressed high levels of *HOPX*, which is commonly used as an AT1 cell marker but has been shown recently to be expressed by multiple epithelial subpopulations in murine lungs [48].

To determine whether the relative abundance of specific epithelial subpopulations was associated with developmental ALV/BM we performed a correlation analysis between the epithelial clusters and sample-level ALV/BM scores (**Figure 4E**). This identified a significant positive correlation for “AT1” and “AT2” clusters with ALV; and showed the basal subpopulation to correlate with BM.

Cell-of-origin is a key determinant of NSCLC subtype [45,49]. However, histological transformation can occur in response to therapy [45]. Given the varied cell-of-origin for LUAD and LUSC, we hypothesised that the mechanisms of BM activation would vary between subtypes: LUSC solely requiring the expansion of malignant basal cells; and LUAD requiring proliferative expansion and de-differentiation from an AT2 to basal phenotype. To investigate this, we first confirmed that BM activation and basal cell abundance correlated after excluding LUSC samples (r = 0.64, p = 2.01e-07, **Figure S4B**). We then utilised the HLCA dataset as a reference to perform label transfer analysis from normal AT2 cells to the epithelial cells present in our dataset using the probability of these classifications as a measure of AT2 lineage fidelity (**Figure 4F**). As expected, cells in our AT2 cluster (from both normal and LUAD tissues) had very high lineage fidelity (median > 0.95); whereas both inflamed and basal cells from LUAD tissues had almost ubiquitously lost AT2 lineage fidelity (**Figure 4F-G**). We then performed a similar analysis of basal lineage fidelity (**Figure 4H**). This showed that the Basal LUAD cluster had only limited fidelity to the basal lineage (median = 0.18), suggesting most of these cells have acquired an atypical basal-like state rather than undergoing complete transdifferentiation to a basal cell phenotype (**Figure 4H-I**). Notably, the Inflamed LUAD population had minimal basal lineage fidelity (median = 0; **Figure 4I**).

To examine the phenotype of LUAD-basal cells further we performed differential gene expression analysis comparing LUAD cells from Inflamed and Basal clusters (**Figure 4J-K**). This showed LUAD-basal cells downregulated expression of alveolar and club cell marker genes (e.g., *SFTPD*, *SFTPB*, *SFTPA2*, *ABCA3*, *NAPSA*, *SCGB3A2*). LUAD-basal cells upregulated *S100A9*; certain genes also highly expressed by LUSC basal cells (*e.g. KRT17*, *KRT19*); and proliferation-associated genes (*e.g. MKI67, TOP2A*). However, canonical basal cell markers (e.g., *KRT14*, *KRT5*, *TP63 and DAPL1*) were only detected in a small proportion of LUAD-basal cells and were not found to be significantly upregulated (**Figure S4C**).

We then stratified LUAD-basal cells into two groups based on the Basal Lineage Fidelity score (“typical” > 0.5 and “atypical” < 0.5): 69% of LUAD-basal cells were classified as atypical and 31% as typical basal cells. We found that the typical LUAD basal cells expressed higher levels of canonical basal markers such as *KRT5, KRT6A,* and *KRT17* (**Figure 4L**), as well as lower levels of alveolar markers, including *SFTPB, NAPSA,* and *ABCA3,* compared to the atypical LUAD basal cells (**Figure S4D**).

To summarise, in LUAD tumours differentiated alveolar cells are lost and replaced predominantly by malignant cells with an inflamed or basal-like phenotype. In cases with BM activation the malignant LUAD cells were characterised by an increased abundance of basal-like cells. These cells upregulated genes highly expressed by LUSC; but to a lower level, suggesting that LUAD-basal cells represented a distinct phenotype rather than a complete histological transformation.

### BM activation is found in high grade LUAD tumours

The association between BM activation and loss of alveolar differentiation prompted us to test whether ALV/BM signatures were linked to morphological grade. Using the TCGA cohort we found that high grade (poorly differentiated) LUAD tumours have significantly higher BM scores than low grade tumours (**Figure 5A**), which was also confirmed in an independent LUAD cohort (**Figure S5A**). In contrast, no significant differences were found in ALV/BM scores for LUSC tumours of different grades (**Figure S5B**).

**Figure 5.**
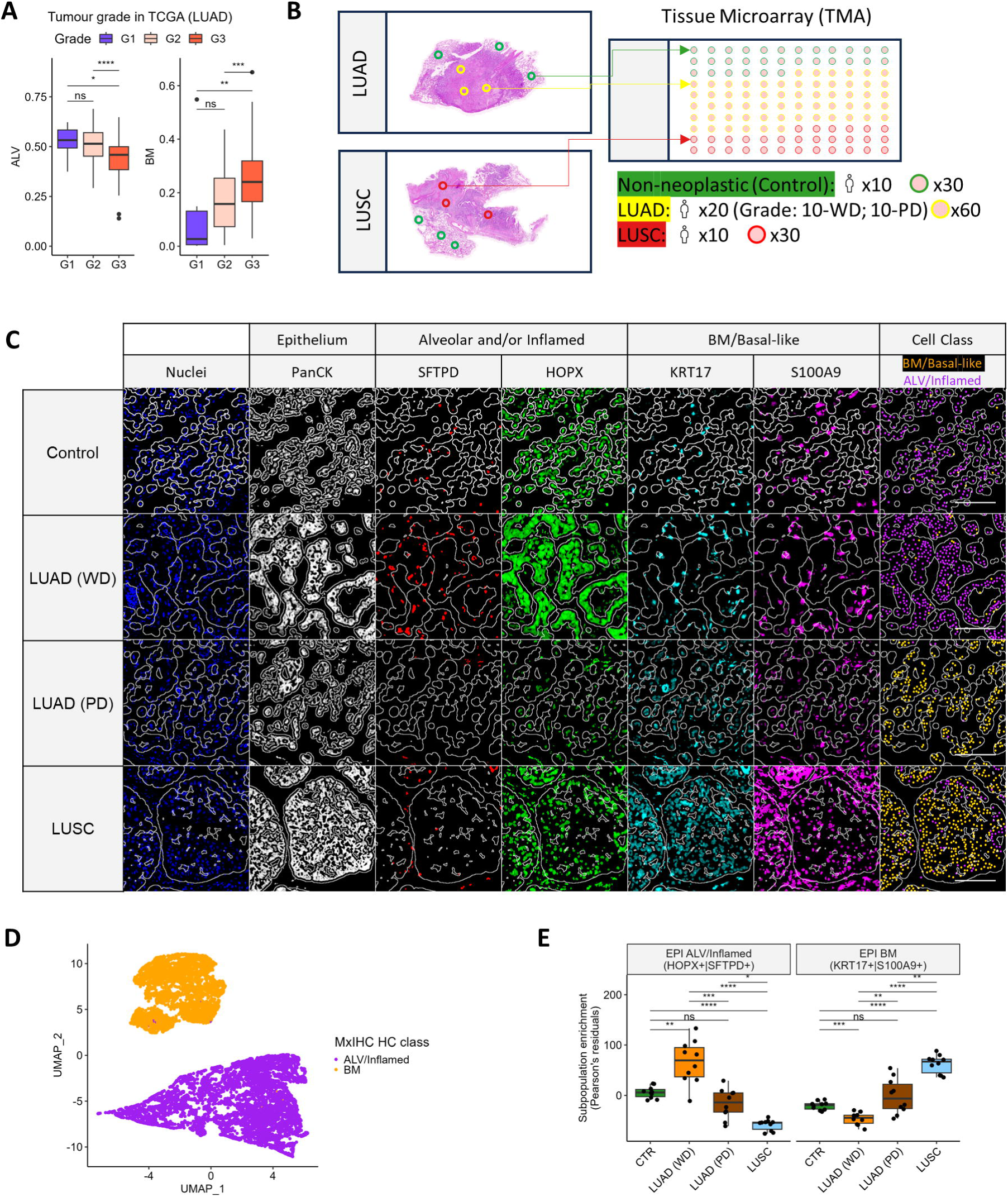
BM activation is found in high grade LUAD tumours. **A** – Boxplots for LUAD grade and ALV/BM scores (TCGA; G1 - well differentiated, G2 - moderately differentiated, G3 - poorly differentiated). **B** – Graphical representation of tissue sampling from LUAD and LUSC blocks to form a tissue microarray (TMA) used for multiplexed Immunohistochemistry (WD – well differentiated; PD – poorly differentiated [solid morphology]). **C** – Representative images from multiplexed immunohistochemistry (mxIHC) analysis of NSCLC TMAs showing pseudo immunofluorescence (pseudoIF; blue staining represents nuclei) and cell classification from histo-cytometry analysis (Cell class) of epithelial (PanCK+) cell expression of BM markers (KRT17, S100A9) and ALV/Inflamed markers (SFTPD, HOPX). White lines indicate the outline of Pan-CK+ tissue regions. **D** – UMAP of histo-cytometry measurements from mxIHC analysis showing HC class assignments, plot displays a randomly sampled 5% of epithelial cells to minimise over-plotting. **E** – Enrichment of Epi ALV/Inflamed and Epi-BM subpopulations across NSCLC samples measured by mxIHC. Wilcoxon rank sum test with Bonferroni correction used to calculate p values in A, E.

To examine the relationship between tumour morphology and the molecular profile of epithelial cells at the protein level, we utilised multiplexed immunohistochemistry (mxIHC) analysis of tissue microarrays (TMAs). For this analysis we curated a cohort of archival FFPE samples to incorporate control (non-neoplastic) lung tissue; low grade (well differentiated [WD]) LUAD; high grade (poorly differentiated [PD] - solid morphology) LUAD; and LUSC (10 samples per group, 3 cores per sample; **Figure 5B**). MxIHC was performed using an antibody panel designed to identify epithelial cells (Pan-Cytokeratin [PanCK]+) expressing BM markers (S100A9 and KRT17) or ALV/inflamed markers (SFTPD and HOPX) (**Figure 5C**). In addition, MKI67 and SOX9 were examined as phenotypic markers of proliferation and developmental BM, respectively. UMAP analysis showed that the BM+ and ALV/inflamed+ subpopulations were clearly discriminated using this panel (**Figure 5D** and **Figure S5C**). Consistent with our transcriptomic analysis, BM+ cells were increased in LUSC compared to LUAD; and in poorly differentiated LUAD compared to well-differentiated LUAD (**Figure 5E**). Furthermore, MKI67 expression was significantly increased in BM+ cells compared to ALV/Inflamed+ cells across all tumour sample subtypes (**Figure S5D**); whereas SOX9 expression was similarly expressed in both ALV/Inflamed+ and BM+ cells from well differentiated LUAD but significantly increased in BM+ compared to ALV/inflamed+ cells from poorly differentiated LUAD or LUSC (**Figure S5E**).

## Discussion

We have demonstrated that developmental programmes (ALV and BM) frequently become dysregulated in NSCLC, with BM activation identifying aggressive LUAD that were resistant to multiple therapies. We found that BM activation in LUAD was associated with *TP53* pathway mutations and required AT2 cells to lose their alveolar identity, acquiring a basal-like state. These findings highlight an important mechanism of disease progression and therapy resistance in LUAD.

BM activation in LUAD was associated with resistance-to and recurrence-from multiple therapeutic strategies, including TKIs and ICB. TKI resistance is known to be in part driven by on-target mutations, such as EGFR T790M mutations, but the mechanisms of resistance in other cases are poorly understood. Our results suggest that BM activation is a critical determinant of TKI resistance and likely underpins the association previously described between TP53 mutations and TKI resistance [50], providing an opportunity for refined biomarker development. Similarly, our results suggest that assessing BM activation could improve upon the current gold standard ICB response biomarkers (TMB and PDL1 expression [51]), which are known to be sub-optimal. Furthermore, the consistent association of BM activation with poor survival and therapy resistance highlights this process as a fundamental mechanism of disease progression, presenting opportunities to identify novel therapeutic targets.

Loss of AT2 lineage fidelity has been described as an early event in KRAS driven LUAD [52]; representing a key factor in disease progression and TKI therapy resistance [28]. We showed that BM activation involved concomitant loss of AT2 lineage fidelity and transdifferentiation to a basal-like phenotype. Basal cell phenotypes have been linked to aggressive subtypes of breast, pancreatic and bladder cancers: associated with stemness [53], invasion [54], metastatic dissemination [55] and therapy resistance [56]. Notably our analysis showed BM activation was strongly linked to TP53 pathway mutations; but reduced in cases with *KRAS* mutations and wild-type *TP53*. During murine lung development hyperactive Kras was shown to be sufficient to induce Sox9 expression, leading to extensive branching and suppressed alveolar differentiation [9]. In murine models of Kras-driven LUAD, p53’s tumour suppressive properties include restricting cellular plasticity and promoting AT1-like differentiation [7,57]. Similarly, in bleomycin induced fibrosis p53 activity is required for regeneration of AT1 cells from AT2 progenitors, with loss of p53 function causing AT2-cells to stall in a transient state marked by *Krt19* expression [58]. This transient state shared many features with KRT5^-^KRT17^+^ basaloid cells found in human IPF state, consistent with the basal-like phenotype we identified in human LUAD. Based on this evidence, it is likely that LUAD tumours achieve loss of AT2 lineage fidelity via multiple means, with *KRAS* or *EGFR* mutations representing common examples of genetic drivers, but acquisition of basal-like features requires loss of p53 function. Furthermore, as p53 activity is known to limit stem cell renewal and promote differentiation, these basal-like features are likely linked to cancer stemness [59]. Given our finding that ALV-BM+ cells were the only epithelial population increased in ICB resistant LUAD cases the acquisition of basal-like features in addition to loss of lineage fidelity may be highly relevant in the context of immune evasion.

A notable limitation to our study is the limited number of LUSC samples analysed, which may be why our findings failed to provide significant insight into the mechanisms of progression in this subtype. Future studies should consider expanding this analysis to a larger cohort of LUSC samples.

## Conclusions

In conclusion, we have shown that BM activation in LUAD tumours is a key mechanism of disease progression and therapy resistance with potential utility as a prognostic and predictive biomarker. We have demonstrated that BM activation is driven by malignant cells undergoing loss of AT2 lineage fidelity and acquisition of a basal-like phenotype and that these cells are enriched in both TKI and ICB recurrence. Therefore, suggesting that therapeutic interventions that manipulate these developmental programmes and prevent transdifferentiation to a basal-like state could lead to better clinical management of LUAD patients.

## Declarations

### Ethics approval and consent to participate

Ethical approval for accessing human tissue samples was obtained through the UK National Research Ethics Service (NRES; Rec No. 10/H0504/32), and informed consent was obtained from each patient. All tissue collection and storage were handled by the Department of Cellular Pathology at University Hospital Southampton NHS Trust. In addition, this study was reviewed and approved by the University of Southampton Faculty of Medicine Ethics Committee and Research Integrity and Governance team.

### Availability of data and materials

The datasets used and/or analysed during the current study are available from the corresponding author on reasonable request.

### Authors’ contributions

K. J. B. and C. J. H. contributed to conceptualization, data curation, formal analysis, validation, visualization and writing (original draft); K. J. B., S. G., N. Z., M. E., M. A. L., J. A. and S. P. contributed to investigation; S. J. C., A. A., E. C. S., C. H. O. and G. J. T. contributed to resources; C. H. O., G. J. T. and C. J. H. contributed to funding acquisition; G. J. T. and C. J. H. contributed to supervision and writing (review and editing).

## Acknowledgements

The authors thank the patients involved in this study and the University of Southampton High Performance Computing team for facilitating data processing and analysis. The results presented are in part based upon data generated by the TCGA Research Network: http://cancergenome.nih.gov/.

## References

1. Bray F, Ferlay J, Soerjomataram I, Siegel RL, Torre LA, Jemal A. Global cancer statistics 2018: GLOBOCAN estimates of incidence and mortality worldwide for 36 cancers in 185 countries. CA Cancer J Clin. 2018;68:394– 424.

2. Vokes NI, Pan K, Le X. Efficacy of immunotherapy in oncogene-driven non-small-cell lung cancer. Ther Adv Med Oncol. 2023;15:175883592311614.

3. Herbst RS, Morgensztern D, Boshoff C. The biology and management of non-small cell lung cancer. Nature. 2018;553:446–54.

4. Zappa C, Mousa SA. Non-small cell lung cancer: current treatment and future advances. Transl Lung Cancer Res. 2016;5:288–300.

5. Laughney AM, Hu J, Campbell NR, Bakhoum SF, Setty M, Lavallée V-P, et al. Regenerative lineages and immune-mediated pruning in lung cancer metastasis. Nat Med. 2020;26:259–69.

6. Kotton DN, Morrisey EE. Lung regeneration: mechanisms, applications and emerging stem cell populations. Nat Med. 2014;20:822–32.

7. Marjanovic ND, Hofree M, Chan JE, Canner D, Wu K, Trakala M, et al. Emergence of a High-Plasticity Cell State during Lung Cancer Evolution. Cancer Cell. 2020;38:229–246.e13.

8. Ferone G, Song J-Y, Sutherland KD, Bhaskaran R, Monkhorst K, Lambooij J-P, et al. SOX2 Is the Determining Oncogenic Switch in Promoting Lung Squamous Cell Carcinoma from Different Cells of Origin. Cancer Cell. 2016;30:519–32.

9. Chang DR, Alanis DM, Miller RK, Ji H, Akiyama H, McCrea PD, et al. Lung epithelial branching program antagonizes alveolar differentiation. Proc Natl Acad Sci U S A. 2013;110:18042–51.

10. Schittny JC. Development of the lung. Cell Tissue Res. 2017;367:427–44.

11. Wu T, Hu E, Xu S, Chen M, Guo P, Dai Z, et al. clusterProfiler 4.0: A universal enrichment tool for interpreting omics data. The Innovation. 2021;2:100141.

12. Barbie DA, Tamayo P, Boehm JS, Kim SY, Moody SE, Dunn IF, et al. Systematic RNA interference reveals that oncogenic KRAS-driven cancers require TBK1. Nature. 2009;462:108–12.

13. Hänzelmann S, Castelo R, Guinney J. GSVA: Gene set variation analysis for microarray and RNA-Seq data. BMC Bioinformatics. 2013;14.

14. Colaprico A, Silva TC, Olsen C, Garofano L, Cava C, Garolini D, et al. TCGAbiolinks: an R/Bioconductor package for integrative analysis of TCGA data. Nucleic Acids Res. 2016;44:e71–e71.

15. Love MI, Huber W, Anders S. Moderated estimation of fold change and dispersion for RNA-seq data with DESeq2. Genome Biol. 2014;15:550.

16. Bakr S, Gevaert O, Echegaray S, Ayers K, Zhou M, Shafiq M, et al. A radiogenomic dataset of non-small cell lung cancer. Sci Data. 2018;5:180202.

17. Wilks C, Zheng SC, Chen FY, Charles R, Solomon B, Ling JP, et al. recount3: summaries and queries for large-scale RNA-seq expression and splicing. Genome Biol. 2021;22:323.

18. Robinson MD, McCarthy DJ, Smyth GK. edgeR: a Bioconductor package for differential expression analysis of digital gene expression data. Bioinformatics. 2010;26:139–40.

19. Lin J, Marquardt G, Mullapudi N, Wang T, Han W, Shi M, et al. Lung Cancer Transcriptomes Refined with Laser Capture Microdissection. Am J Pathol. 2014;184:2868–84.

20. Schabath MB, Welsh EA, Fulp WJ, Chen L, Teer JK, Thompson ZJ, et al. Differential association of STK11 and TP53 with KRAS mutation-associated gene expression, proliferation and immune surveillance in lung adenocarcinoma. Oncogene. 2016;35:3209–16.

21. Okayama H, Kohno T, Ishii Y, Shimada Y, Shiraishi K, Iwakawa R, et al. Identification of genes upregulated in ALK-positive and EGFR/KRAS/ALK-negative lung adenocarcinomas. Cancer Res. 2012;72:100–11.

22. Shedden K, Taylor JMG, Enkemann SA, Tsao M-S, Jurisica I, Giordano TJ, et al. Gene expression–based survival prediction in lung adenocarcinoma: a multi-site, blinded validation study. Nat Med [Internet]. 2008;14:822–7. Available from: http://www.nature.com/doifinder/10.1038/nm.1790%5Cnpapers3://publication/doi/10.1038/nm.1790

23. Bueno R, Richards WG, Harpole DH, Ballman K V., Tsao MS, Chen Z, et al. Multi-Institutional Prospective Validation of Prognostic mRNA Signatures in Early Stage Squamous Lung Cancer (Alliance). Journal of Thoracic Oncology. 2020;15:1748–57.

24. Raponi M, Zhang Y, Yu J, Chen G, Lee G, Taylor JMG, et al. Gene Expression Signatures for Predicting Prognosis of Squamous Cell and Adenocarcinomas of the Lung. Cancer Res [Internet]. 2006;66:7466–72. Available from: http://cancerres.aacrjournals.org/lookup/doi/10.1158/0008-5472.CAN-06-1191

25. ME R, B P, D W, Y H, CW L, W S, et al. Limma powers differential expression analyses for RNA-sequencing and microarray studies. Nucleic Acids Res. 2015;43(7):e47.

26. Kassambara A, Mundt F. Factoextra: Extract and Visualize the Results of Multivariate Data Analyses. R package. 2020;

27. Kim N, Kim HK, Lee K, Hong Y, Cho JH, Choi JW, et al. Single-cell RNA sequencing demonstrates the molecular and cellular reprogramming of metastatic lung adenocarcinoma. Nat Commun. 2020;11.

28. Maynard A, McCoach CE, Rotow JK, Harris L, Haderk F, Kerr DL, et al. Therapy-Induced Evolution of Human Lung Cancer Revealed by Single-Cell RNA Sequencing. Cell. 2020;182:1232–1251.e22.

29. Zilionis R, Engblom C, Pfirschke C, Savova V, Zemmour D, Saatcioglu HD, et al. Single-Cell Transcriptomics of Human and Mouse Lung Cancers Reveals Conserved Myeloid Populations across Individuals and Species. Immunity [Internet]. 2019;50:1317–1334.e10. Available from: https://linkinghub.elsevier.com/retrieve/pii/S1074761319301268

30. Hanley CJ, Waise S, Parker R, Lopez MA, Taylor J, Kimbley L, et al. Spatially discrete signalling niches regulate fibroblast heterogeneity in human lung cancer. bioRxiv [Internet]. 2020;2020.06.08.134270. Available from: https://www.biorxiv.org/content/10.1101/2020.06.08.134270v1

31. Travaglini KJ, Nabhan AN, Penland L, Sinha R, Gillich A, Sit R V., et al. A molecular cell atlas of the human lung from single-cell RNA sequencing. Nature [Internet]. 2020;587:619–25. Available from: 10.1038/s41586-020-2922-4

32. Satija R, Farrell JA, Gennert D, Schier AF, Regev A. Spatial reconstruction of single-cell gene expression data. Nat Biotechnol. 2015;33:495–502.

33. Stuart T, Butler A, Hoffman P, Hafemeister C, Papalexi E, Mauck WM, et al. Comprehensive Integration of Single-Cell Data. Cell. 2019;177:1888–1902.e21.

34. Hu J, Zhang L, Xia H, Yan Y, Zhu X, Sun F, et al. Tumor microenvironment remodeling after neoadjuvant immunotherapy in non-small cell lung cancer revealed by single-cell RNA sequencing. Genome Med. 2023;15:14.

35. Mayakonda A, Lin D-C, Assenov Y, Plass C, Koeffler HP. Maftools: efficient and comprehensive analysis of somatic variants in cancer. Genome Res. 2018;28:1747–56.

36. Gaujoux R, Seoighe C. A flexible R package for nonnegative matrix factorization. BMC Bioinformatics. 2010;11:367.

37. Hanley CJ, Waise S, Ellis MJ, Lopez MA, Pun WY, Taylor J, et al. Single-cell analysis reveals prognostic fibroblast subpopulations linked to molecular and immunological subtypes of lung cancer. Nat Commun. 2023;14:387.

38. Mammoto A, Mammoto T. Vascular Niche in Lung Alveolar Development, Homeostasis, and Regeneration. Front Bioeng Biotechnol. 2019;7.

39. Wang X, Ricciuti B, Nguyen T, Li X, Rabin MS, Awad MM, et al. Association between Smoking History and Tumor Mutation Burden in Advanced Non–Small Cell Lung Cancer. Cancer Res. 2021;81:2566–73.

40. Chua KP, Teng YHF, Tan AC, Takano A, Alvarez JJS, Nahar R, et al. Integrative Profiling of T790M-Negative EGFR-Mutated NSCLC Reveals Pervasive Lineage Transition and Therapeutic Opportunities. Clinical Cancer Research. 2021;27:5939–50.

41. Jung H, Kim HS, Kim JY, Sun J-M, Ahn JS, Ahn M-J, et al. DNA methylation loss promotes immune evasion of tumours with high mutation and copy number load. Nat Commun. 2019;10:4278.

42. Lin C, Shi X, Zhao J, He Q, Fan Y, Xu W, et al. Tumor Mutation Burden Correlates With Efficacy of Chemotherapy/Targeted Therapy in Advanced Non–Small Cell Lung Cancer. Front Oncol. 2020;10.

43. Assoun S, Theou-Anton N, Nguenang M, Cazes A, Danel C, Abbar B, et al. Association of TP53 mutations with response and longer survival under immune checkpoint inhibitors in advanced non-small-cell lung cancer. Lung Cancer. 2019;132:65–71.

44. Negrao M V., Lam VK, Reuben A, Rubin ML, Landry LL, Roarty EB, et al. PD-L1 Expression, Tumor Mutational Burden, and Cancer Gene Mutations Are Stronger Predictors of Benefit from Immune Checkpoint Blockade than HLA Class I Genotype in Non–Small Cell Lung Cancer. Journal of Thoracic Oncology. 2019;14:1021–31.

45. Sato Y, Saito G, Fujimoto D. Histologic transformation in lung cancer: when one door shuts, another opens. Ther Adv Med Oncol. 2022;14:175883592211305.

46. LaFave LM, Kartha VK, Ma S, Meli K, Del Priore I, Lareau C, et al. Epigenomic State Transitions Characterize Tumor Progression in Mouse Lung Adenocarcinoma. Cancer Cell. 2020;38:212–228.e13.

47. Chan M, Liu Y. Function of epithelial stem cell in the repair of alveolar injury. Stem Cell Res Ther. 2022;13:170.

48. Liu K, Meng X, Liu Z, Tang M, Lv Z, Huang X, et al. Tracing the origin of alveolar stem cells in lung repair and regeneration. Cell. 2024;187:2428–2445.e20.

49. Wang Z, Li Z, Zhou K, Wang C, Jiang L, Zhang L, et al. Deciphering cell lineage specification of human lung adenocarcinoma with single-cell RNA sequencing. Nat Commun. 2021;12:6500.

50. Ferrara MG, Belluomini L, Smimmo A, Sposito M, Avancini A, Giannarelli D, et al. Meta-analysis of the prognostic impact of TP53 co-mutations in EGFR-mutant advanced non-small-cell lung cancer treated with tyrosine kinase inhibitors. Crit Rev Oncol Hematol. 2023;184:103929.

51. Li X, Song W, Shao C, Shi Y, Han W. Emerging predictors of the response to the blockade of immune checkpoints in cancer therapy. Cell Mol Immunol. 2019;16:28–39.

52. Dost AFM, Moye AL, Vedaie M, Tran LM, Fung E, Heinze D, et al. Organoids Model Transcriptional Hallmarks of Oncogenic KRAS Activation in Lung Epithelial Progenitor Cells. Cell Stem Cell. 2020;27:663–678.e8.

53. Papafotiou G, Paraskevopoulou V, Vasilaki E, Kanaki Z, Paschalidis N, Klinakis A. KRT14 marks a subpopulation of bladder basal cells with pivotal role in regeneration and tumorigenesis. Nat Commun. 2016;7:11914.

54. Cheung KJ, Gabrielson E, Werb Z, Ewald AJ. Collective Invasion in Breast Cancer Requires a Conserved Basal Epithelial Program. Cell. 2013;155:1639– 51.

55. Cheung KJ, Padmanaban V, Silvestri V, Schipper K, Cohen JD, Fairchild AN, et al. Polyclonal breast cancer metastases arise from collective dissemination of keratin 14-expressing tumor cell clusters. Proceedings of the National Academy of Sciences. 2016;113.

56. Ruta V, Naro C, Pieraccioli M, Leccese A, Archibugi L, Cesari E, et al. An alternative splicing signature defines the basal-like phenotype and predicts worse clinical outcome in pancreatic cancer. Cell Rep Med. 2024;5:101411.

57. Kaiser AM, Gatto A, Hanson KJ, Zhao RL, Raj N, Ozawa MG, et al. p53 governs an AT1 differentiation programme in lung cancer suppression. Nature. 2023;619:851–9.

58. Kobayashi Y, Tata A, Konkimalla A, Katsura H, Lee RF, Ou J, et al. Persistence of a regeneration-associated, transitional alveolar epithelial cell state in pulmonary fibrosis. Nat Cell Biol. 2020;22:934–46.

59. Garrido-Jimenez S, Barrera-Lopez JF, Diaz-Chamorro S, Mateos-Quiros CM, Rodriguez-Blanco I, Marquez-Perez FL, et al. p53 regulation by MDM2 contributes to self-renewal and differentiation of basal stem cells in mouse and human airway epithelium. The FASEB Journal. 2021;35.

