## Supplementary material for "Developmental programmes drive cellular plasticity, disease progression and therapy resistance in lung adenocarcinoma": Supplemantary Figures and Tables: Supplementary Figures.pdf

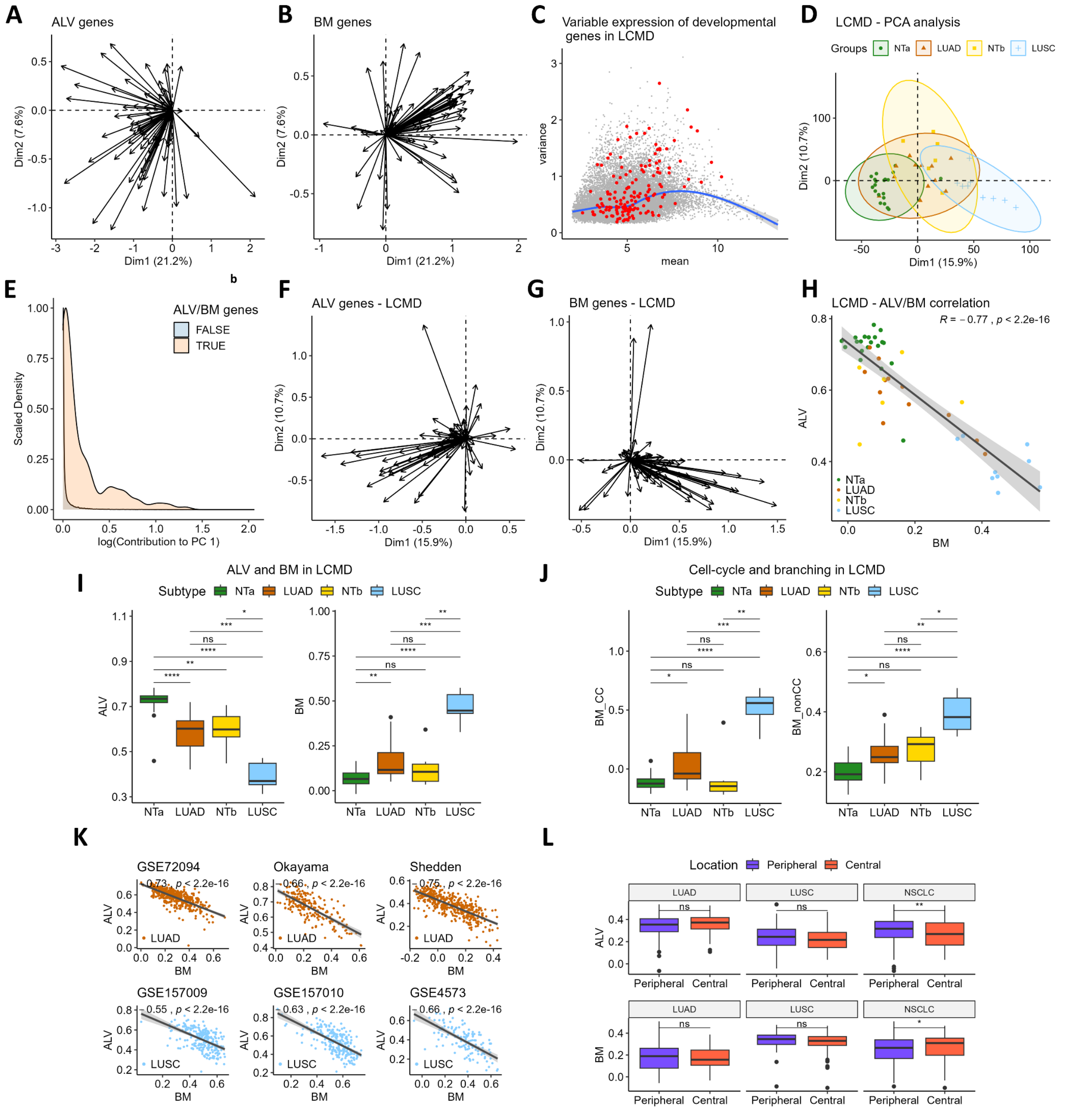


**Figure S1**. **Developmental Alveogenesis (ALV) and Branching Morphogenesis (BM) programmes are associated with transcriptomic variance in NSCLC. A/B** – PCA loading plots for ALV genes (**A**) and BM genes (**B**) in TCGA RNA-Seq data. **C** – Scatter plot showing mean expression levels and variance for all genes in NSCLC samples from Laser-Capture-Microdissection-Dataset (LCMD) [1]. BM and ALV genes are shown in red. **D-G** – PCA of NSCLC samples from LCMD. **D** – PC plot showing sample separation. **E** – Histogram showing the contribution of ALV/BM genes PC1 (LCMD). **F,G** - PCA loading plots for ALV genes (**F**) and BM genes (**G**) in LCMD. **H** – Scatter plot showing ssGSEA scores for ALV/BM programmes across LCMD samples. R = Spearman's correlation. **I,J**- Comparison of enrichment of ALV/BM (**I**) as well as BM_cc (cell-cycle related genes) and BM_nonCC (non-cell-cycle related genes) (**J**) across LCMD NSCLC samples. NTa – non-tumour alveolar, Ntb – non-tumour bronchial. **K** – Scatter plot showing ALV/BM scores in LUAD/LUSC microarray datasets. LUAD: GSE72094 [2], Okayama (GSE31210) [3], Shedden

(GSE68465) [4]. LUSC: GSE157009 (cohort I) [5], GSE157010 (cohort II) [5], GSE4573 [6]. R = Spearman’s correlation. **L** - Comparison of ssGSEA scores for ALV/BM programmes between LUAD tumours, LUSC tumours, and NSCLC tumours (LUAD and LUSC combined) located in either peripheral or central parts of the lung (TCGA). P values calculated with Wilcoxon test (I,J,L) and Bonferroni adjusted (I,J).

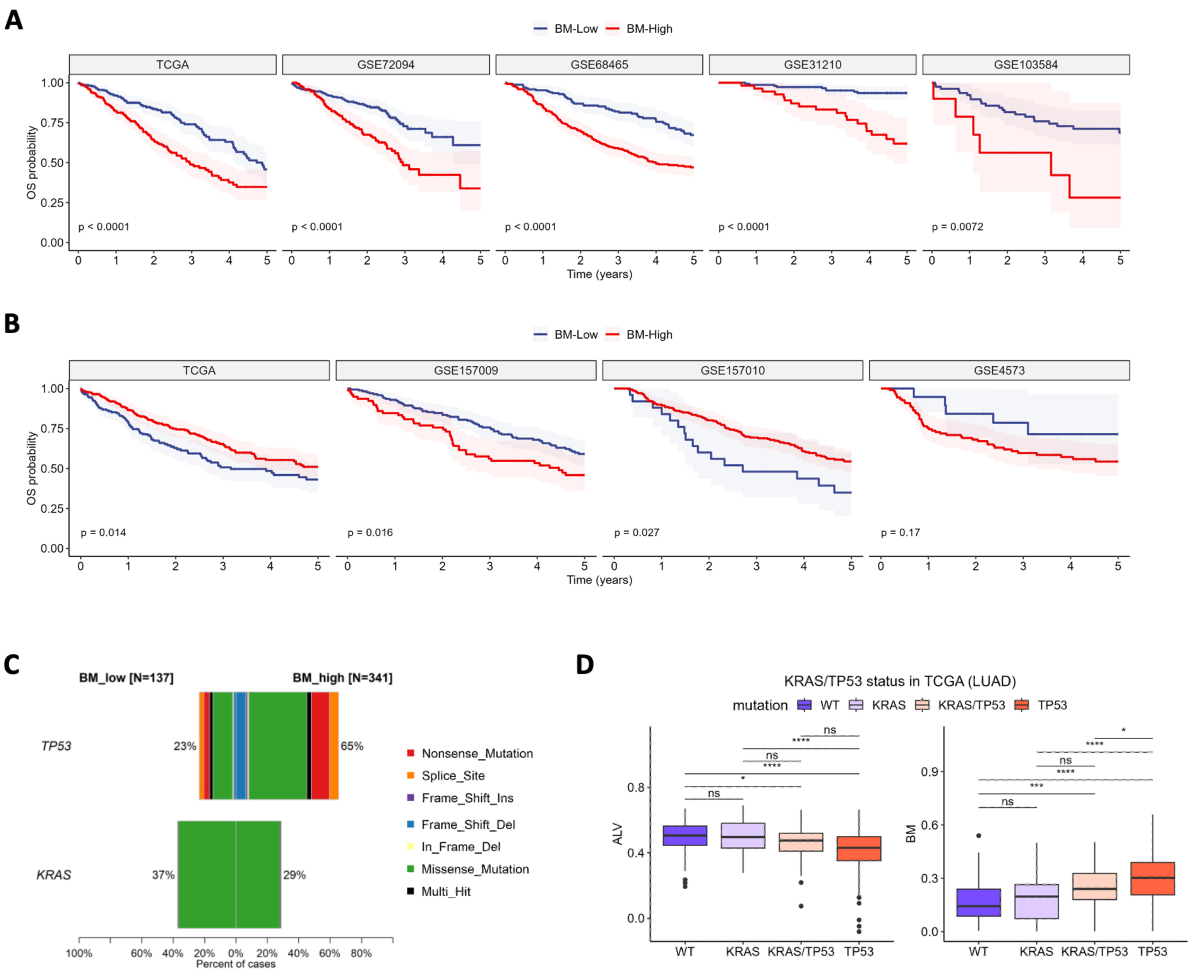


**Figure S2. High expression of BM in LUAD predicts poor survival and is associated with frequent *TP53* mutations. A** – KM plots showing 5-year overall survival (OS) based on BM expression across 5 LUAD datasets: TCGA, GSE72094 [2], GSE68465 (Shedden) [4], GSE31210 (Okayama) [3], GSE103584 [3]. P values calculated with log-rank test; shaded region represents 95% CIs. **B** - KM plots showing 5-year overall survival (OS) based on BM expression across 4 LUSC datasets: TCGA, GSE157009 [5], GSE157010 [5], GSE4573 [6]. **C** – Percentage of LUAD TCGA samples with TP53 or KRAS mutations. **D** – Comparison of expression of ALV/BM across LUAD TCGA samples with KRAS mutations (WT, TP53), TP53 mutations (WT, KRAS), both KRAS and TP53 mutations, or neither of the two (WT).

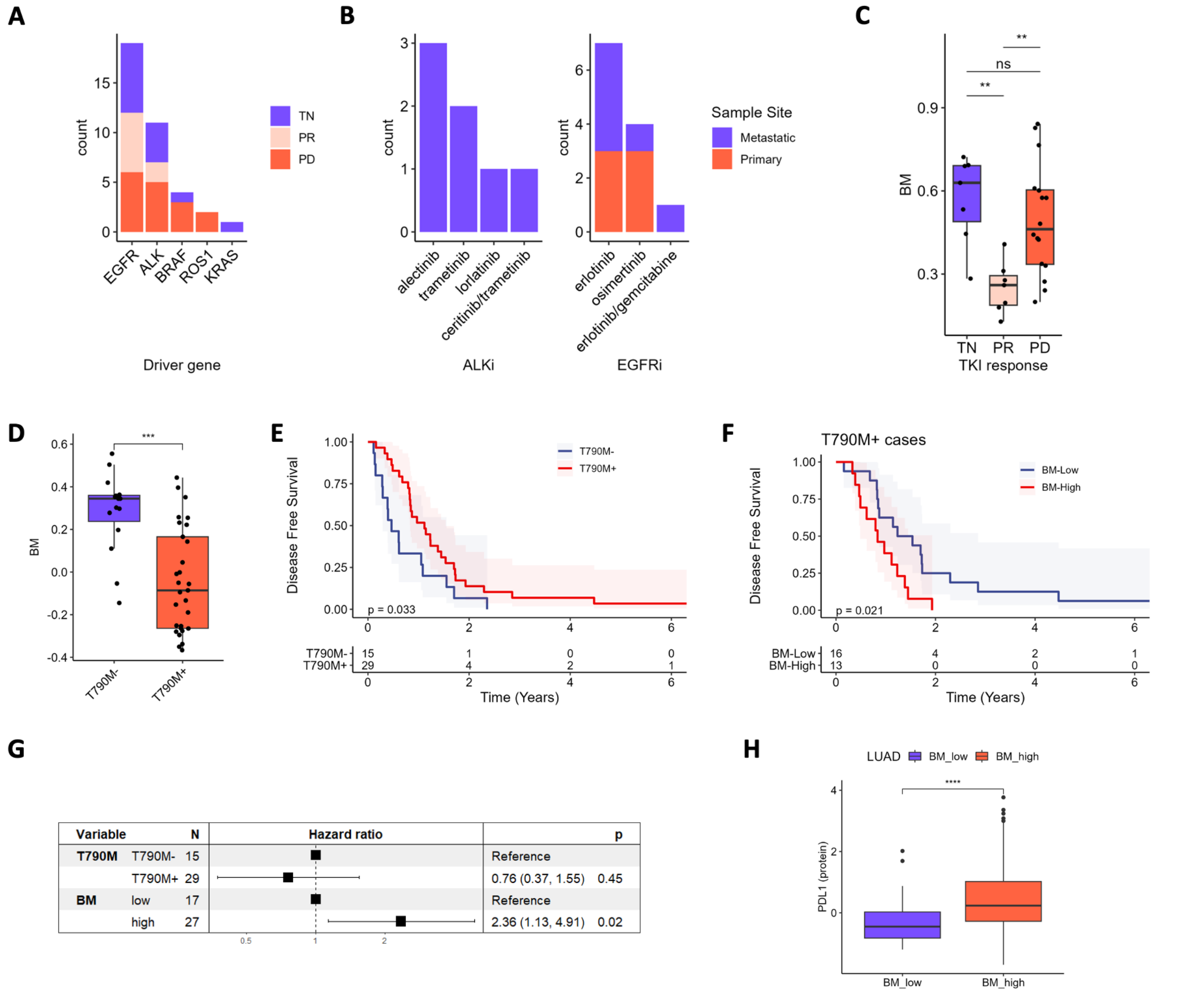


**Figure S3. High expression of BM is associated with resistance to therapies**. **A** – Barplot showing the number of samples analysed form the Maynard cohort [7] with the indicated driver gene and their associated grouping for analysis; TN – treatment naïve, PR – partial response, PD – progressive disease. **B** – Barplots showing the TKI treatment received for post-treatment samples with ALK or EGFR drivers. **C** – Boxplot showing the BM score for Stage IV samples only from the Maynard cohort. **D** – Boxplot showing BM scores in the IMPACT (Chua) [8] cohort grouped by T790M status.   number of samples rec of post-treatment samples. **E** – Kaplan-Meier plot showing disease free survival rates for the IMPACT cohort stratifying patients by T790M status. F – Kaplan-Meier plot showing disease free survival rates for T790M+ cases in the IMPACT cohort stratified by BM status. **G** – Forest plot showing multivariate regression analysis of the IMPACT cohort including T790M and BM status as independent variables. **H** - RPPA (reverse phase protein array) data of PD-L1 expression in LUAD TCGA samples grouped by BM expression. Asterisks represent P values calculated with Wilcoxon test with Bonferroni correction where multiple tests were performed.

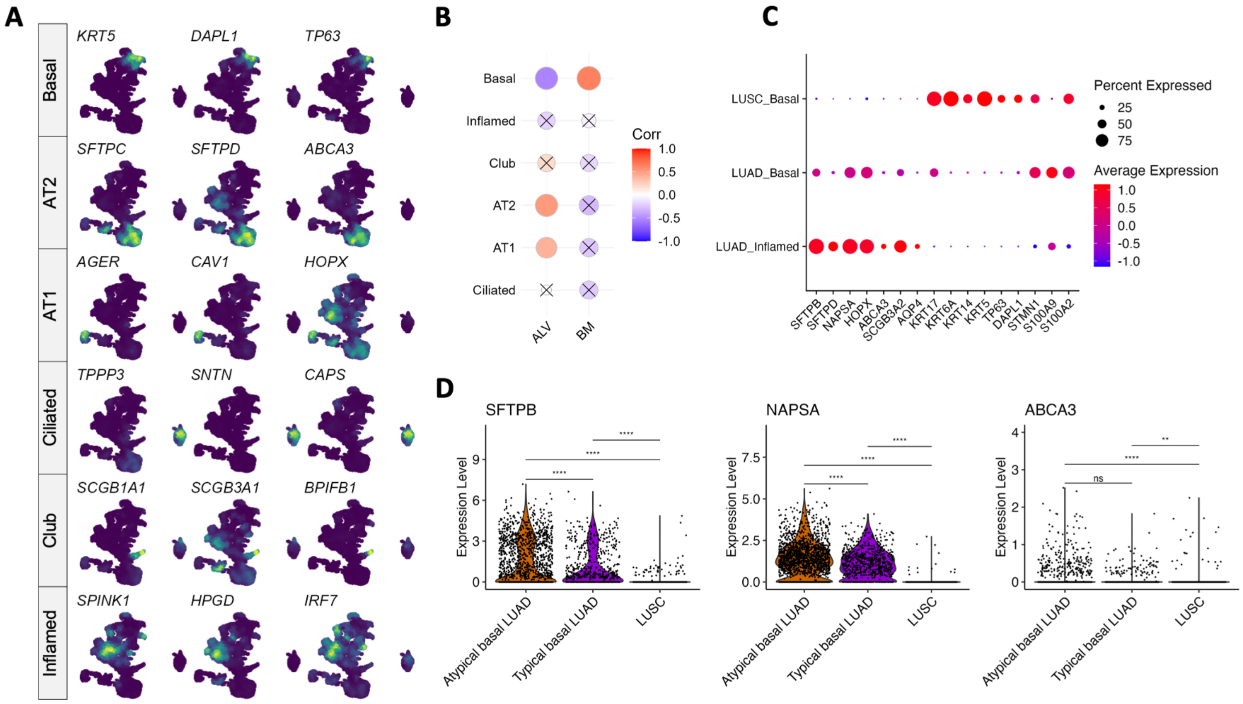


**Figure S4. BM upregulation in LUAD is associated with acquisition of basal-like features. A** – Feature density plots showing example marker genes for epithelial subpopulations. **B** - Sample-level Pearson's correlation between ssGSEA scores for ALV/BM signatures and epithelial cluster abundance after excluding LUSC samples. P values > 0.01 are crossed out. **C –** Dot plot showing expression of alveolar and basal associated genes in LUAD-Inflamed, LUAD-Basal and LUSC-Basal cells. **D** – Violin plots showing expression of alveolar genes in LUAD and LUSC cells from the Basal cluster. LUAD cells with a basal prediction > 0.5 = "Typical basal cells"; < 0.5 = "Atypical basal cells". Wilcoxon rank sum test with Bonferroni's correction.

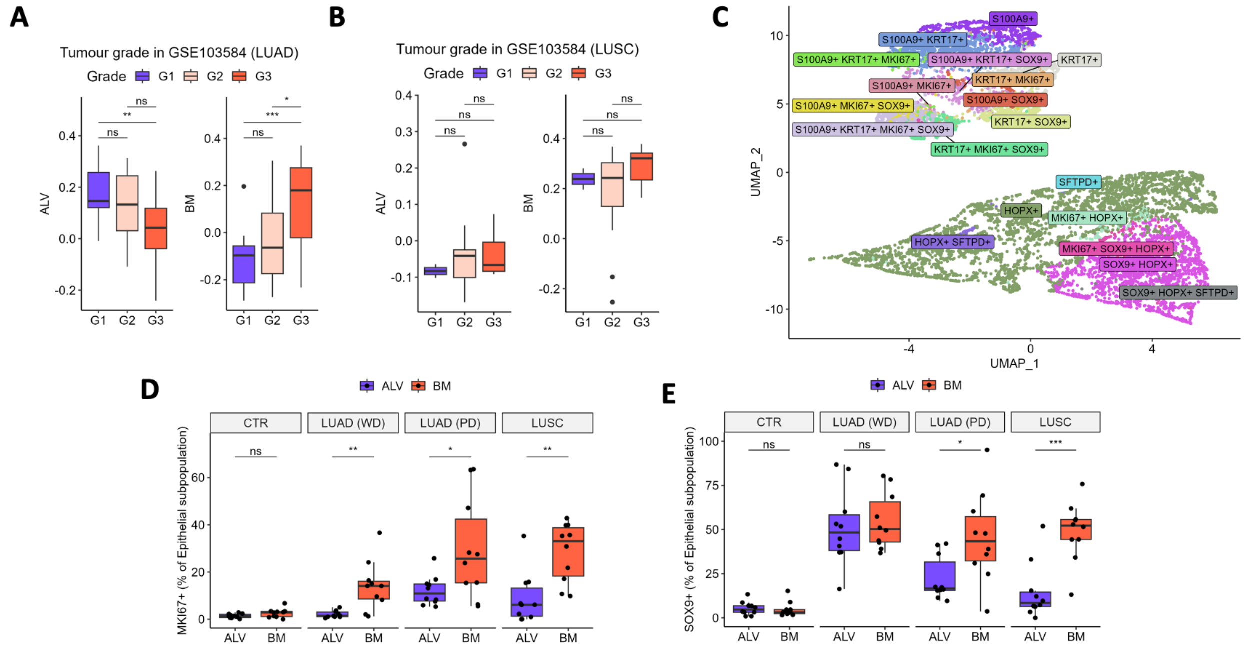


**Figure S5. BM in LUAD is associated with high grade tumors. A-B** - Association between tumour grade and BM score in LUAD (**A**) and LUSC (**B**) (GSE103584) [9] samples. G1 - well differentiated, G2 - moderately differentiated, G3 - poorly differentiated. **C** – UMAP representation of NSCLC mxIHC epithelial (PANCK+) cells assigned to ALV/Inflamed or BM phenotypes, coloured by the complete immune-phenotype. **D-E** - Boxplot showing the difference in the percentage of MKI67-positive (**D**) and SOX9-positive epithelial cells (**E**) between BM population (S100A9+/KRT17+) and ALV/Inflamed population (SFTPD+/HOPX+) across NSCLC mxIHC samples. Asterisks represent P values calculated with Wilcoxon test with Bonferroni correction where multiple tests were performed.
